# *rnaends*: an R package to study exact RNA ends at nucleotide resolution

**DOI:** 10.1101/2025.04.18.649472

**Authors:** Tomas Caetano, Peter Redder, Gwennaele Fichant, Roland Barriot

## Abstract

5’ and 3’ RNA-end sequencing protocols have unlocked new opportunities to study aspects of RNA metabolism such as synthesis, maturation and degradation, by enabling the quantification of exact ends of RNA molecules *in vivo*. From RNA-Seq data that have been generated with one of the specialized protocols, it is possible to identify transcription start sites (TSS) and/or endoribonucleolytic cleavage sites, and even, in some cases, co-translational 5’ to 3’ degradation dynamics. Furthermore, post-transcriptional addition of ribonucleotides at the 3’ end of RNA can be studied at the nucleotide resolution.

While different RNA-end sequencing library protocols exist that have been adapted to a specific organism (prokaryote or eukaryote) or specific biological question, the generated RNA-Seq data are very similar and share common processing steps. Most importantly, the major aspect of RNA-end sequencing is that only the 5’ or 3’ end mapped location is of interest, contrary to conventional RNA sequencing that considers genomic ranges for gene expression analysis. This translates to a simple representation of the quantitative data as a count matrix of RNA-end location on the reference sequences. This representation seems under-exploited and is, to our knowledge, not available in a generic package focused on the analyses on the exact transcriptome ends.

Here, we present the *rnaends* R package which is dedicated to RNA-end sequencing analysis. It offers functions for raw read pre-processing, RNA-end mapping and quantification, RNA-end count matrix post-processing, and further downstream count matrix analyses such as TSS identification, fast Fourier transform for signal periodic pattern analysis, or differential proportion of RNA-end analysis. The use of *rnaends* is illustrated here with applications in RNA metabolism studies through selected *rnaends* workflows on published RNA-end datasets: (i) TSS identification, (ii) ribosome translation speed and co-translational degradation, (iii) post-transcriptional modification analysis and differential proportion analysis.

## Introduction

Since its first development, high throughput sequencing has been derived into numerous applications in molecular biology, notably for genomics and post-genomics studies. The preparation of sequencing libraries for short read sequencing generally involves fragmentation of the polynucleotide prior to ligation of sequencing adapters. For transcriptome studies, this fragmentation renders the identification of the exact 5’ and 3’ termini of RNA molecules challenging at the nucleotide resolution. Knowledge about the exact ends of RNAs opens various opportunities in the study of RNA metabolism. On the 5’ end, (i) it can determine the exact transcription start site (TSS) positions (and consequently also promoter positions) as in (Prados et al., 2016), or (ii) it can give access to 5’ end degradation intermediates revealing *in vivo* footprint of ribosome occupancy and dynamics (Pelechano et al., 2015; Nersisyan et al., 2020), or (iii) it can locate endoribonucleolytic cleavage sites (Khemici et al., 2015; Broglia et al., 2020). On the 3’ end, it can locate endoribonucleolytic cleavage sites and transcription termination sites (TTSs) (Broglia et al., 2020), and it is also able to identify (and quantify the abundance of) ribonucleotides added post-transcriptionally to RNAs (Xu et al., 2023). To avoid the loss of precision due to RNA fragmentation, specialised RNA-Seq library preparation protocols have been developed to obtain reads that correspond only to the exact 5’ ends or 3’ ends of RNA molecules (Sharma et al., 2010; Sharma & Vogel, 2014).

In eukaryotes, for the mRNA 5’ end identification, many protocols for library preparation rely on the cap protecting the 5’ end of messenger RNA from degradation by exoribonuclease, or, for the 3’ end studies, on the polyA tail. To analyse such specific data, dedicated software and tools have been developed. For the identification of 5’ ends, the CAGE (Murata et al., 2014) and GRO-cap (Core et al., 2014) protocols are recommended (Adiconis et al., 2018) and, for downstream analyses, packages such as *CAGEr* (Haberle et al., 2015) and *CAGEfightR* (Thodberg et al., 2019) are readily available.

In prokaryotes, newly synthesized RNAs harbour a tri-phosphate at their 5’ end and a hydroxyl at their 3’ end (denoted respectively 5’PPP and 3’OH hereafter). Several protocols adapted for these molecules have been developed for RNA-Seq library preparation such as EMOTE (Redder, 2015), PARE (DiChiara et al., 2016), and Term-seq (Dar et al., 2016). The common principle of these protocols is to ligate a specific adapter (a specific RNA or DNA oligonucleotide) to the 5’ or 3’ termini of RNA molecules as illustrated in Figure 1. Ligation only occurs for RNA ends that exhibit specific biochemical properties such as a mono-phosphate at the 5’ end (5’P) or a hydroxyl at the 3’ end (3’OH). After ligation of this specific adapter, the RNA is reverse transcribed to cDNA, whereupon cDNA generated from RNAs ligated to the adapter are amplified by PCR. This is accomplished by having one of the PCR primers hybridising to an invariable sequence of the ligated oligo (we refer to this invariable sequence as the “recognition sequence”). Apart from the recognition sequence, the ligated oligo frequently contains additional features to help downstream analyses, such as a random stretch of nucleotides to serve as a unique molecular identifier (UMI) to eliminate PCR amplification bias, and a control sequence to unambiguously pinpoint the junction between oligo and RNA.

**Figure 1.**
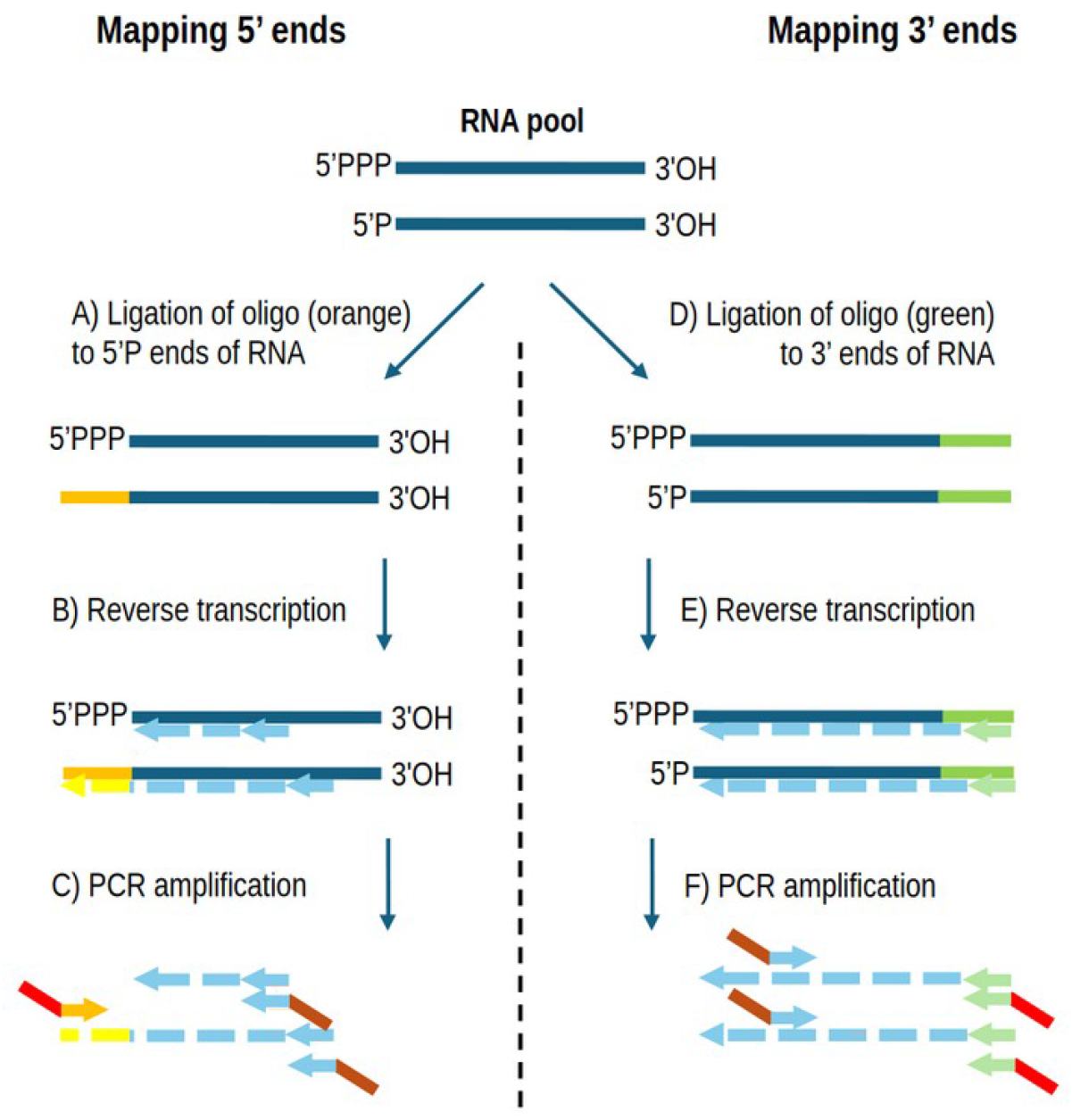
Key steps in RNA-Seq library preparation protocols for determining the exact RNA-ends. Details vary according to protocol used. Mapping 5’ ends (panels A, B and C): A) Ligation of an oligo with known sequence (orange) to 5’P ends of RNA. RNAs with 5’PPP are not accepted as substrate for the ligase, but if needed, then the RNA can be pre-treated with a pyrophosphohydrolase to convert 5’PPP ends to 5’P (Sharma & Vogel, 2014; Prados et al., 2016). B) Reverse transcription to generate cDNA (light blue). For ligated RNAs, the cDNA will terminate with the reverse complement of the oligo sequence (yellow). C) PCR to amplify the cDNA and add adapter sequences for NGS sequencing (red and brown). Only cDNAs from ligated RNAs will be amplified, since the red/orange primer can only hybridise to the yellow sequence (thus the upper RNA in the panel will not be amplified and will not include the red adaptor). Sequencing is performed starting from the red adapter sequence. Mapping 3’ ends (panels D, E and F): D) Ligation of an oligo with known sequence (green) to 3’OH ends of RNA. E) Reverse transcription to generate cDNA (light blue) that will start with the reverse complement of the oligo sequence (light green). Only ligated RNAs will be reverse transcribed. F) PCR to amplify the cDNA and add adapter sequences for NGS sequencing (red and brown). Only cDNA from ligated RNAs will be amplified. Sequencing is performed starting from the red adapter sequence.

The exact protocol used for the RNA-Seq library preparation will vary depending on the issue addressed by the study of RNA metabolism. However, the sequencing data generated are very similar and share common processing steps to quantify RNA-end abundances at the nucleotide resolution. Indeed, once the reads are aligned to the reference genome, only the 5’-most or the 3’-most mapped position is of interest for further analyses. Therefore, unlike conventional high-throughput sequencing analyses, that generally involve taking into account genomic ranges corresponding to mapped reads when quantifying RNAs, RNA-end sequencing quantification only considers reads that map to the same 5’-most or 3’-most position. This reduced, but powerful, representation is, to our knowledge, not readily available and exploited in existing software. Therefore, we are faced with the absence of a generic package focused on the analyses of the exact ends of transcriptomes when studying RNA metabolism. Indeed, in the previously published studies (*e.g*. (Sharma et al., 2010); (Prados et al., 2016); (Dar et al., 2016); (Xu et al., 2023)), each research group has developed their own tools and scripts to pre-process and analyse the same type of data to address biological questions related to RNA metabolism.

Over the years, while studying the RNA maturation and decay in *Staphylococcus aureus*, we have developed basic tools as well as more elaborate functions that we have integrated in an R package named *rnaends* presented here. The main principles driving this package are (i) pre-processing of reads prior to their alignment to a reference sequence, (ii) quantification of mapped reads on reference sequences at single nucleotide locations (5’ end or 3’ end) which is given in an output count table, (iii) downstream studies performed by statistical analyses available in the R environment or in a Bioconductor package, or an *ad hoc* implementation provided by our package.

In the following, we present the key *rnaends* R package features that we have organized into different topics corresponding to different phases of typical analysis projects such as those depicted in Figure 2. We first describe the functions developed for the pre-processing of sequencing reads to validate and convert raw sequencing reads into reads mappable to reference sequences (Figure 2B, 2C and 2D). Then, we describe how mapped reads are quantified to obtain a count table representing RNA-end abundances at the nucleotide resolution (Figure 2E and 2F). Finally, we illustrate the use of the *rnaends* package for the investigation of selected questions related to the study of RNA metabolism. Starting with raw read data available from publications, we describe the workflows implemented from read pre-processing to statistical analyses applied to the resulting count table.

**Figure 2.**
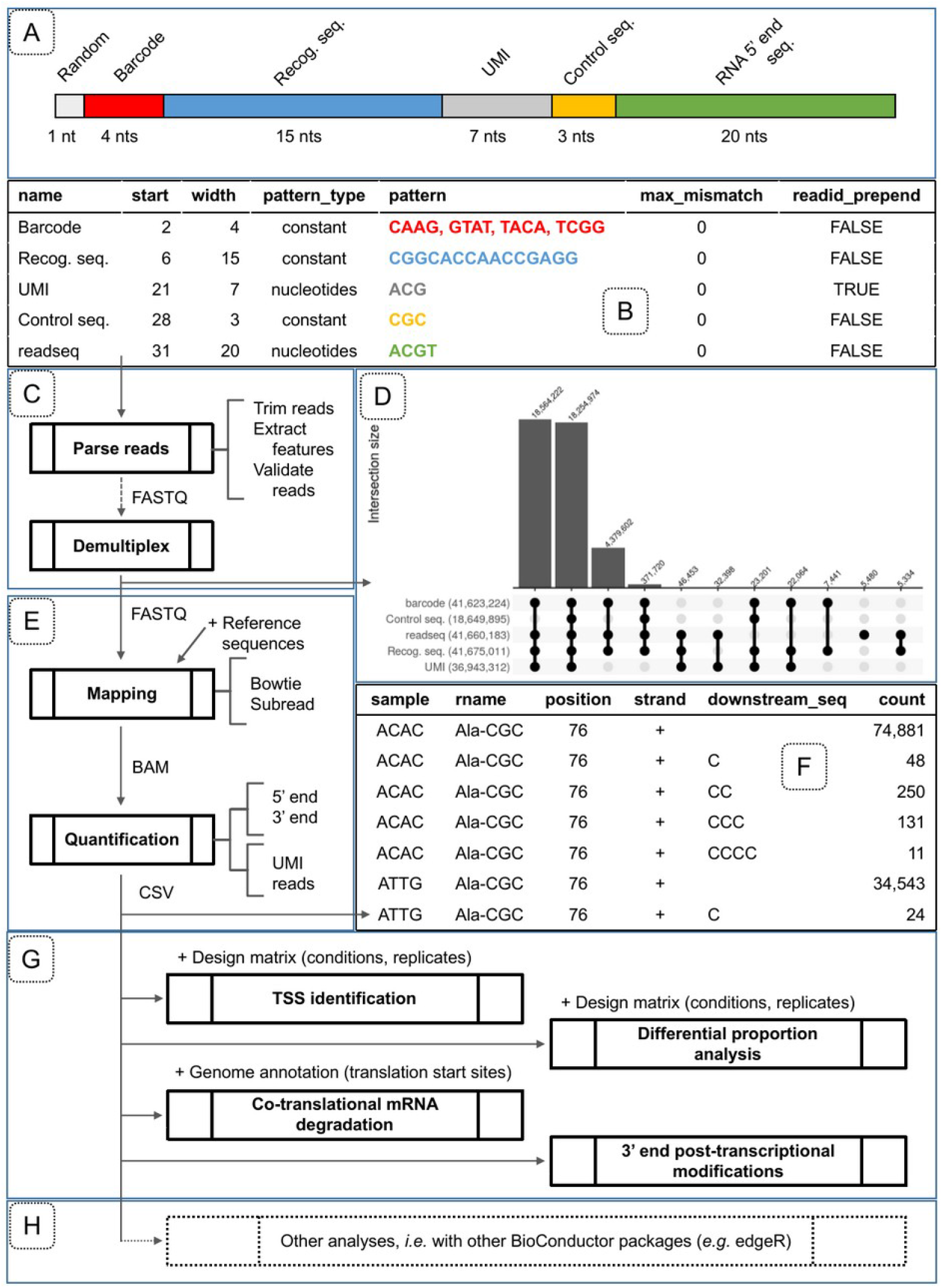
Overview of the features provided in the *rnaends* package grouped by topics of an RNA-end sequencing project. A) Overview of an example of expected read structure of 5’ end sequencing: 1 random nucleotide, then the 4 nucleotides of barcode (here either CAAG, GTAT, TACA or TCGG) identifying 4 multiplexed samples, then the recognition sequence (CGGCACCAACCGAGG in this example), then a UMI (7 random A, C or G nucleotides in this example), then 3 nucleotides of control sequence (CGC in this example), and eventually the actual 5’ end of the RNA (20 nucleotides in this example, but will typically be longer when examining eukaryotes) which can be mapped onto the reference sequences. The length of each element can be defined by the user, according to the experimental setup and the examined organism); B) Example of a *read_features* table obtained with the functions *init_read_features* and *add_read_feature* corresponding to the read structure in panel A. max_mismatch: The number of allowed mismatches when checking for the presence of the feature. readid_prepend: TRUE if the sequence of the feature should be kept associated with the read for subsequent analysis steps; C) FASTQ reads pre-processing and validation based on a *read_features* table describing the structure of sequencing reads; D) Distribution of read features validity described in panel A and represented by an upset plot generated after the parsing, validation and demultiplexing of a raw read FASTQ file. Each combination of valid features is displayed in the lower upset plot while the upper bar plot shows the corresponding number of reads. This plot is highly useful for determining whether the wet-lab experiment worked well, and if not, then where the problem arose; E) Reads alignment on reference sequences and abundance computation at the single nucleotide resolution to obtain: F) a count table representing the abundances (*count*) of RNA 3’ ends mapped to a reference sequence (*rname*) at a certain location (*position*) on a *strand* and with the indicated post-transcriptionally added nucleotide sequence (*downstream_seq*) that could not be aligned to the reference sequence. In this example of 3’ end sequencing (from Xu et al 2023), the reads contain UMI extracted in the pre-processing step (with readid_prepend set to TRUE) that are used at this step to remove PCR amplification duplicates introduced in the library preparation; G) *rnaends* provides functions for downstream analysis of the count table for TSS identification, analysis of RNA-end species proportions (*e.g*. 5’P/5’PPP in WT *vs*. mutant strain), periodic signal analysis such as ribosome translation speed profiling, or the exploration of 3’ end modifications. The user provides a design matrix to link each count to a given biological replicate and the associated experimental conditions. H) *rnaends* count table can also be analysed by other packages.

## Material and methods

### Development of *rnaends* as an R package

*rnaends* is implemented in R version 4.4.2. The choice of using only R and R packages available from CRAN or Bioconductor was motivated by the availability of downstream statistical analyses and the broad spectrum of packages offered by the Bioconductor ecosystem (Huber et al., 2015). The aim of the package is also to provide non expert users with an entry point for RNA-end sequencing projects by providing simple but versatile functions for processing FASTQ files and computing 5’ or 3’ end count table.

The main functions provided in the package are listed in Table 1, grouped by analysis-topic and are detailed in the next subsections. For the pre-processing of reads and the quantification of exact RNA-end abundances steps, we rely on widely used Bioconductor packages that provide FASTQ and BAM files processing to compute the count table. Afterwards, we make use of the *tidyverse* metapackage to further process the count table and integrate some downstream analyses. The choice of *tidyverse* was motivated by the extensive documentation available on-line, and the wide functionalities available for processing and formatting tables (with the included packages *tidyr, dplyr, tibbles)*, as well as their visualisation (with the *ggplot2* package).

**Table 1.** Main *rnaends* package functions grouped by RNA-end sequencing study steps indicated between parentheses: (α) specification of the sequencing read structure expected in a FASTQ file as a *read_features* table (example in Figure 2B), and its use for pre-processing sequencing reads; (β) reads alignment to reference sequences and quantification of the exact 5’ or 3’ ends as a count table; (γ) downstream analyses of the count table for different types of applications integrated in the package.

| Function (study step) | Description |
| --- | --- |
| init_read_features ($\alpha$) | Initializes and returns a read_features table with the minimal feature (named readseq) corresponding to the region in the read sequence that is expected to align to a reference sequence |
| add_read_feature ($\alpha$) | Adds a feature to the read_features table returned by init_read_features |
| parse_reads ($\alpha$) | Parses each read from a FASTQ file to retrieve features, test their validity and extract sequence to align |
| demultiplex ($\alpha$) | Demultiplexes a FASTQ file according to barcode sequence reads provided through a read_features object |
| map_fastq ($\beta$) | Performs the mapping of reads of a FASTQ file using either bowtie or subread aligner with appropriate parameters for specifying the number of mismatches allowed in the alignments (default: max_map_mismatch=0) and if matches should be unique (default: map_unique=TRUE). |
| quantify_5prime ($\beta$) | Converts aligned reads on reference sequences from a BAM file to the count table of exact 5' ends |
| quantify_3prime ($\beta$) | Converts aligned reads on reference sequences from a BAM file to the count table of exact 3' ends and unmapped downstream sequences |
| TSS_betabinomial ($\gamma$) | Computes the p-value that a specific genomic location (position) is a TSS, using a beta-binomial model. |
| TSS ($\gamma$) | Performs identification of Transcription Start Sites (TSS) over all positions from a 5' end experiment |
| differential_proportion ($\gamma$) | Performs identification of significant differences in proportions over all positions from a 5' or 3' end experiment. For example for a given TSS, the proportion that is of monophosphorylated. |
| metacounts_normalized ($\gamma$) | Aggregates and normalizes the counts of individual positions within defined genomic regions. Counts are normalized within each region to equalize the effect of highly versus less transcribed regions. |
| metacounts_feature ($\gamma$) | Computes metacounts for a window around a set of locations of a given feature (e.g. 1<sup>st</sup> nucleotide of all translation start codons) |
| metacounts_region ($\gamma$) | Computes metacounts for a set of regions (e.g. start locations and widths of all CDS) |
| FFT ($\gamma$) | Formats counts or metacounts for one or multiple regions and calls FFT_profile to performs signal periodicity analysis |
| FFT_profile ($\gamma$) | Performs fast Fourier transform signal periodicity analysis on a count or metacount profile |
| frame_preference ($\gamma$) | Computes global translation frame preference over a set of regions (e.g. all CDS) from the exact 5' end count table. See section "Ribosome translation speed and co-translational degradation" |

The *rnaends* functions are provided in an R package distributed *via* the GitLab platform. Source code is available as a git repository on GitLab at https://gitlab.com/rnaends/rnaends and documentation and tutorial workflows are available on GitLab Pages at https://rnaends.gitlab.io/rnaends (see Data, scripts, code, and supplementary information availability section).

### Hardware requirements and performances

Running times and memory used were benchmarked on a Dell Latitude 5280 laptop with dual core CPU Intel i5-7300 with 16GB memory. For a typical use case, 6 samples of 40 million sequencing reads each, multiplexed in a FASTQ file of 240 millions of reads, the pre-processing step takes ∼2.5 hours (∼30k reads/sec) and less than 1.5GB of memory. The mapping to reference sequences takes 1.5 hour and 500MB of memory. The computation of the count matrix takes ∼45 minutes and ∼8GB of memory. Then, downstream analysis of the count matrix (TSS identification), takes 2 minutes and 1GB of memory.

### Data

Datasets used in this manuscript and the R notebooks in the on-line documentation make use of publicly available datasets accompanying publications from others (see Data, scripts, code and supplementary information availability section for more details).

For the co-translational degradation analysis, the RNA-Seq datasets used from (Hou et al., 2016) are PARE libraries (Zhai et al., 2014) obtained on inflorescences of *Arabidopsis thaliana*. The library preparation protocol is as follows. Polyadenylated RNAs are isolated. From this pool, only 5’P RNAs are ligated (i) with an adapter, and reverse transcribed using an oligo(dT) with a 3’ adapter sequence. Second strand synthesis and PCR amplification are performed. Then, *MmeI* digestion generates signature fragments 20 bp downstream of 5’P RNAs ligated in (i). Digestion products are ligated with a double-stranded 3’-DNA adapter. After PCR amplification and PAGE-purification, the sequencing library is generated.

For the post-transcriptional 3’ end addition analysis, the four RNA-Seq datasets used from (Xu et al., 2023) were multiplexed in an RNA-Seq library with other samples. The library preparation protocol is as follows. tRNAs were *in vitro* transcribed from full length tRNA templates harbouring CCA at their 3’ end. From these samples, four samples were prepared: two incubated with the MenT1 toxin and two without the toxin. Then samples were ligated at their 3’OH end with a synthetic oligo harbouring an UMI sequence of length 15 followed by a reverse transcription step that incorporates a barcode for multiplexing samples for sequencing.

### Reads parsing, validation, trimming, feature extraction and demultiplexing

As illustrated in Figure 1, RNA-end sequencing library preparation generally relies on the ligation of a synthetic oligonucleotide to RNAs. Consequently, the ligated oligo sequence will be present in the raw sequencing reads that, therefore, will need to be pre-processed and reformatted prior to their alignment on the reference sequences. In *rnaends*, sequencing reads are expected to be stored in the FASTQ file format, and are processed with the *ShortRead* (Morgan et al., 2009) Bioconductor package version 1.64.0.

Reads corresponding to RNA samples from different conditions or strains might be multiplexed by incorporating an index into the adapter for sequencing. Moreover, further multiplexing is sometimes accomplished by incorporating barcodes into the oligos used for RNA ligation or the PCR primers used for amplification, and/or these oligos may contain functional sequences such as a unique molecule identifier (UMI) to eliminate duplicates resulting from a PCR amplification step that follows the adapter ligation. All these features will be part of the sequencing read itself, and will need to be removed before mapping the read to a reference sequence.

For different published datasets, we identified cases of read pre-processing scenarios and implemented a set of functions to check read validity, to extract features and to format the reads in order to generate FASTQ files ready for the mapping step. To this end, the user must fill up the *read_features* table (illustrated in Figure 2B) to describe the structure and content of the raw sequencing reads (expected pattern type and location on reads). As processing more complex patterns, such as regular expressions, can be time-consuming, *rnaends* uses more specific patterns to achieve better performances on millions of reads. For instance, patterns can consist of (i) a constant string expected at a certain location on the reads (such as a control sequence to validate that the sequence of a ligated oligo is part of the reads), (ii) a set of allowed constant strings (typically barcodes when multiple samples are multiplexed for sequencing), (iii) a random sequence of allowed nucleotides (typically corresponding to the sequence to be aligned to the reference and/or the UMIs). The UMI sequences is an example of a feature for which the sequence should be kept associated with the read, using readid_prepend=TRUE, since the UMI sequence can be used to eliminate PCR bias from the dataset (see section “Mapping and quantification”).

Once the expected read features are set up, the user can pre-process the raw reads FASTQ file to perform the feature matching and extraction, and the reads validation (Figure 2A, 2B and 2C). One or more FASTQ files will be obtained containing only valid reads ready to be aligned to the reference sequences, together with a report containing statistics (with number of reads matching each feature) for each sample on the content and quality of the FASTQ file of raw reads.

For an illustration, we used the dataset provided in (Khemici et al., 2015) where the sequencing reads are 50 nucleotides long and structured as follows: the first sequence nucleotide is a random nucleotide followed by a barcode of 4 nucleotides (CAAG, GTAT, TACA or TCGG) identifying 4 multiplexed samples, followed by a recognition sequence (CGGCACCAACCGAGG), then by a UMI (7 random A, C or G nucleotides), and a control sequence (CGC), and finally by the actual 5’ end of the RNA which is 20 nucleotides long. Figure 2D is an illustration generated with our package on the results obtained from the report, which contains the presence or absence of each valid feature for each read. For the dataset analysed in Figure 2D, a large number of reads are rejected due to lack of control sequence (leftmost bar), suggesting that the forward PCR oligo (red/orange arrow in Figure 1C) was not sufficiently specific for the oligo sequence at the temperature used in the PCR. This information is highly useful to the user who constructed the RNA-Seq library or performed the experiment, allowing future improvements. Then, almost as many reads carry all the valid features (second bar from the left) and will be retained to be aligned to the reference sequence(s). Other invalid reads are due to an invalid UMI and control sequence, or to an invalid UMI only. The rest of the invalid features, or their combination, represent smaller fractions of all the reads.

### Mapping and quantification

The originality of RNA-end sequencing lies in the fact that only the first (respectively last) mapped position is of interest, as it corresponds to the exact 5’ end (respectively 3’ end) of the RNA molecule that was ligated to the adapter during library preparation.

For the read alignment, we have integrated the use of *bowtie* (Langmead et al., 2009) and *subread* (Liao et al., 2013) aligners with default appropriate parameters for the following reasons. Sequencing reads are aligned to reference sequences expected in the FASTA format, and the alignment can be performed to multiple reference sequences if needed (*e.g*. multiple chromosomes or cDNA sequences generated from spliced eukariotic mRNAs). For alignments that should match perfectly, *Rbowtie* (Langmead et al., 2009) version 1.46.0 is used by default with parameters *m=1* (for uniquely mapped reads) and *v=0* (for perfect matches), but these can be adjusted by the user. For sequencing reads that should align partially due to the 3’ end addition of ribonucleotides, *Rsubread* (Liao et al., 2019) version 2.20.0 is used with parameters *unique* and *maxMismatches* set by the user and parameters *output_format=“BAM”, sortReadsByCoordinates=TRUE, type=“dna”, indels=0*. Alignments are produced or converted to BAM files to reduce space.

After mapping, read alignments can be described using a single nucleotide position in the reference sequence, *i.e*. either the 5’ or 3’ end of the RNA. However, in the case where ribonucleotides have been added to the 3’ end after transcription, users should be interested in accessing the sequence absent from the reference sequence. We therefore provide two functions, in the *rnaends* package, one for processing RNA 5’ ends (*quantify_5prime*) and the other one for processing RNA 3’ ends that might possess post-transcriptional modifications (*quantify_3prime)*. Both functions return a count table from a BAM file. In the count table, the mapped reads are counted by reference sequence name (chromosome, cDNA or any sequence provided in the reference sequences FASTA file), position and strand, and in the case of the 3’ end function, the downstream sequence in the read not found in the reference sequence is also provided. An example illustration is given in Figure 2F.

Of note, BAM alignment files can be obtained through the *rnaends* functions but this is not mandatory: users can make use of any software to produce alignments. Alignments to reference sequences are expected in the BAM file format and are processed with *Rsamtools* (Martin Morgan, 2017) version 2.22.0. The *scanBam* function is used to extract mapped read information and converted to count tables stored as *tibbles* of the *tidyverse* meta-package version 2.0.0.

UMIs are most of the time necessary and incorporated in the reads in order to remove PCR amplification or other bias introduced by the library preparation. If UMIs were present in the sequencing reads and specified during the pre-processing step using the *readid_prepend* parameter in the *add_read_feature* function, they can be used in the quantification to filter library preparation amplification biases, *i.e*. the counts can be reported for mapped reads or for unique UMI mapped reads at a location on the reference. It is also possible to keep the UMI sequences included in the count table for further investigation such as their occurrence distribution in case they are not distributed as one could expect from random sequences (this is for example the case in data from (Khemici et al., 2015), data not shown).

As for the read pre-processing step, a statistics report can be generated for each BAM file produced. These statistics include the total number of reads and the percentage of uniquely mapped reads.

### Downstream analyses integrated in the *rnaends* package

Obtaining the exact RNA-end count table from raw sequencing reads is generally the milestone that allows researchers to address the initial questions of their biological study. However through different studies over the years, we have explored and developed several downstream analyses of the exact RNA-end count table that could be useful for other projects. We have therefore integrated them into the *rnaends* package.

#### Differential abundance between paired samples: Transcription Start Site identification

RNA-end sequencing was initially developed to address the inability of RNA sequencing protocols to determine precisely the exact 5’ ends of RNA molecules *in vivo* and their poor performance in identifying and quantifying the preferred transcription start site (TSS) at a genomic location. The strategy in prokaryotes is based on the phosphorylation state of the RNA 5’ end. Indeed, only RNAs which 5’ ends are tri-phosphorylated correspond to native 5’ extremities and therefore to TSSs. An example of a protocol that exploits this is TSS-EMOTE (Prados et al., 2016): Briefly, the library preparation protocol consists in first degrading all 5’P RNAs, and then the RNA is mixed with a synthetic RNA oligo (specific adapter) and split into two pools into which a ligase is added that can only bind the adaptor to 5’P RNAs. One pool serves as control to measure the level of background ligation (noise). The other is treated to convert 5’PPP to 5’P RNAs in order to sequence and quantify all RNA 5’ ends that were initially tri-phosphorylated (see (Prados et al., 2016) for a detailed procedure).

In experimental setups, such as TSS-EMOTE, with two paired samples from the same biological replicate, one with the background distribution and the other with some signal, a simple statistical test relying on a beta-binomial distribution can be used to identify positions for which the signal deviates significantly from the background. This test is implemented in the *TSS_betabinomial* function that takes as input 4 parameters, *i.e*. the total counts in the two samples and the counts at a specific position in the two samples, and returns the *p*-value of the beta-binomial test provided by the *VGAM* package (Yee, 2020) version 1.1. In TSS-EMOTE, biological replicates are needed in the statistical analyses to determine whether a 5’ end is a TSS or not. Thus, *p*-values obtained from replicates are combined with Fisher’s method using the *combine.test* function of *survcomp* (Schröder et al., 2011) version 1.56.0. All positions in an experiment with paired samples and replicates can be tested with the *TSS* function. This function uses the count table accompanied by a design matrix that associates counts to samples and replicates, and returns the *p*-values adjusted for multiple testing with, by default, the False Discovery Rate (Benjamini & Hochberg, 1995).

A tutorial with detailed analysis and code is available in an R notebook provided as supplementary material (see Data, scripts, code, and supplementary information availability section). Given that this approach is simply based on the application of a beta-binomial model between two samples, it can easily be transposed to analyse other experimental designs.

#### Differential proportion between conditions

Certain types of data from analyses of RNA ends are in the form of proportions (*e.g*. the proportion of RNA molecules with 5’P at a given nucleotide with respect to the total number of RNA molecules with a 5’ end at the given nucleotide). When comparing proportions, it is important, from a statistical perspective, to take into account the sequencing depth, *i.e*. the counts for the different species of RNAs in the samples, as well as the replicates, in the different experimental conditions or strains. This leads to counts of instances with a type (*e.g*. 5’P) over the total counts at a specific location, per condition and per replicate. The Bioconductor *DSS* package (Park & Wu, 2016) provides well suited procedures for this type of statistical analysis by estimating the local dispersion on counts from all experimental replicates for a feature (*i.e*. single nucleotide position in our context) and the global dispersion on counts over all features in the sequencing library. The function provided by *DSS* was developed to compute statistics for proportions observed in a DNA region (regardless of the reference genome strand) with counts corresponding to methylated *vs*. all sites (taking replicates into account). In *rnaends* we instead use it on a single nucleotide location on a specific strand. The *DSS* package (Park & Wu, 2016) version 2.54.0 is therefore integrated in the function *differential_proportion* to perform the statistical test of RNA species proportions-difference on a specific location on a given strand. This function uses the count table along with a design matrix that associates counts to samples and replicates, and returns *p*-values and adjusted *p*-values for each strand and nucleotide position.

#### Metacounts profile: aggregate counts of multiple regions to amplify or study a particular feature genome-wide

Periodicity of the ribosomal pauses can be revealed by the RNA 5’ end degradation intermediates quantification (Pelechano et al., 2015). To study the periodicity of a signal around a given feature, such as the start codon of a protein coding gene, the sequencing depth is generally not sufficient, and one might need to compute a metacount profile from all these features on the reference sequences (*i.e*. all start codons) to amplify the signal. The regions from which the counts are to be aggregated can be specified in two ways. First, they can be defined by a fixed window around a set of features (for example 100 residues around the translation start codons), using the *metacounts_feature* function of the package. Second, the set of regions can be specified by their start location and length (for example all the protein coding regions often named CDS), using the *metacounts_region* function. In the latter case, if the regions have different lengths, the length of the metacounts profile is specified by the quantile of the distribution of the region lengths. For instance, a quantile of 0.25 (the default value implying that at least 75% of the regions are longer) will ensure that metacounts at a given position will result from data available in at least 75% of the regions. Smaller regions are filled with zeroes and longer regions are truncated. Once the set of regions of the same length is defined, the core function for metacounts computation is *metacounts_normalized* which extracts and normalizes counts on a set of genome regions. Counts are first normalized by region to avoid biases towards highly expressed regions and varying sequencing depths between samples to be merged. This consists in dividing each count in a region by the total counts of the region, *i.e*. converting them to frequencies. Then, the metacount of a given position in the window is obtained by taking the median of frequencies observed at this position in the set of regions to be combined. By default, the *metacounts_normalized* function returns the metacounts for all the samples and regions combined, but it is also possible to compute them for each feature/region and/or sample, for example when sequencing depth is very large and a strong signal is expected in a region.

#### Signal periodicity

Periodicity in the signal composed of counts in a region or a metacounts profile can be analysed with a fast Fourrier transform analysis. The *rnaends* package provides the *FFT_profile* function which performs the analysis on a single profile with the *fft* function of the *stats* R package. *FFT_profile* returns the fast Fourier transform analysis results formatted from an RNA-end counts analysis perspective as a table in which each row consists in the signal strength detected for a period measured in nucleotides. The *FFT* function of the *rnaends* package provides a wrapper around the *FFT_profile* function to perform signal periodicity analysis over a set of regions, that can be analysed separately or combined as metacounts.

#### Frame preference

The *frame_preference* function of the package is specific to protein coding genes and computes the percentages of counts falling into the three possible translation frame. Counts are normalized over regions through a call to *metacounts_region* prior to their percentage computation.

## Results

### Ribosome translation speed and co-translational degradation

In eukaryotes and in some prokaryotes, degradation of RNA can be achieved by 5’ to 3’ exoribonucleases (Mathy et al., 2007; Hu et al., 2009; Durand et al., 2015). Such degradation of mRNAs can be blocked by translating ribosomes, and the 5’ to 3’ degradation will follow the last translating ribosome (Hu et al., 2009). Quantification of 5’P RNA ends, *i.e*. degradation intermediates, can therefore reveal an *in vivo* footprint of ribosome occupancy and dynamics on translated mRNAs (Pelechano et al., 2015). When 5’ to 3’ degradation is co-translational and at least as fast as translation, ribosomal pauses can typically be observed upstream of the start and stop codons, as well as shorter ribosomal pauses along the protein coding sequences (CDS) resulting in a periodicity of three nucleotides (one codon).

To detect such ribosomal pausing events, we implemented three functions based on the work described in (Nersisyan et al., 2020). These functions make use of the 5’ end quantification accompanied by an annotation table containing start and stop codon locations and potentially other features of interest on reference sequences. We tested our workflow on the raw data published as part of the study of RNA degradation intermediates in *Arabidopsis thaliana*, rice and soybean (Hou et al., 2016). These data were obtained using the PARE protocol (Zhai et al., 2014) in which polyadenylated RNAs are isolated, and a 5’-RNA adapter is ligated only to the RNAs with a 5’P end. As a result only RNAs with a 5’P end will have a 5’ extension that corresponds to the oligonucleotide sequence (see Data section for details).

In short, 5’ end counts can be considered relative to a reference genomic location as the first nucleotide of the start or stop codon, and all 5’ ends that were mapped to a position relative to the chosen location (*e.g*., position −17 upstream of the stop codon) are aggregated over all annotated CDSs to obtain a metacount profile over the user-selected window size around the genomic reference location. This is achieved by applying the *metacounts_feature* function (see Material and methods for details). Figure 3A shows that the positions −16 and −17 relative to the stop codon have the highest values for metacounts of 5’ ends, which could highlight a pause of the ribosome at the translation end site when the ribosome has its A site stalled at a stop codon. Before this pause, it can also be seen that the blue bars corresponding to translation frame 2 have higher values than the other two frames. Periodicity analysis can be achieved by applying a fast Fourier transform (FFT) to the metacounts profile computed on all the CDS. In Figure 3B, the strongest signal is observed for a period of approximately three nucleotides. Furthermore, for a given period such as three nucleotides, metacounts from all CDS can be grouped to obtain a frame preference indication by applying the *frame_preference* function. Figure 3C shows that most of the 5’ end mRNA degradation intermediates in *A. thaliana* align on frame 2 of translation, followed by frame 3, then frame 1.

**Figure 3.**
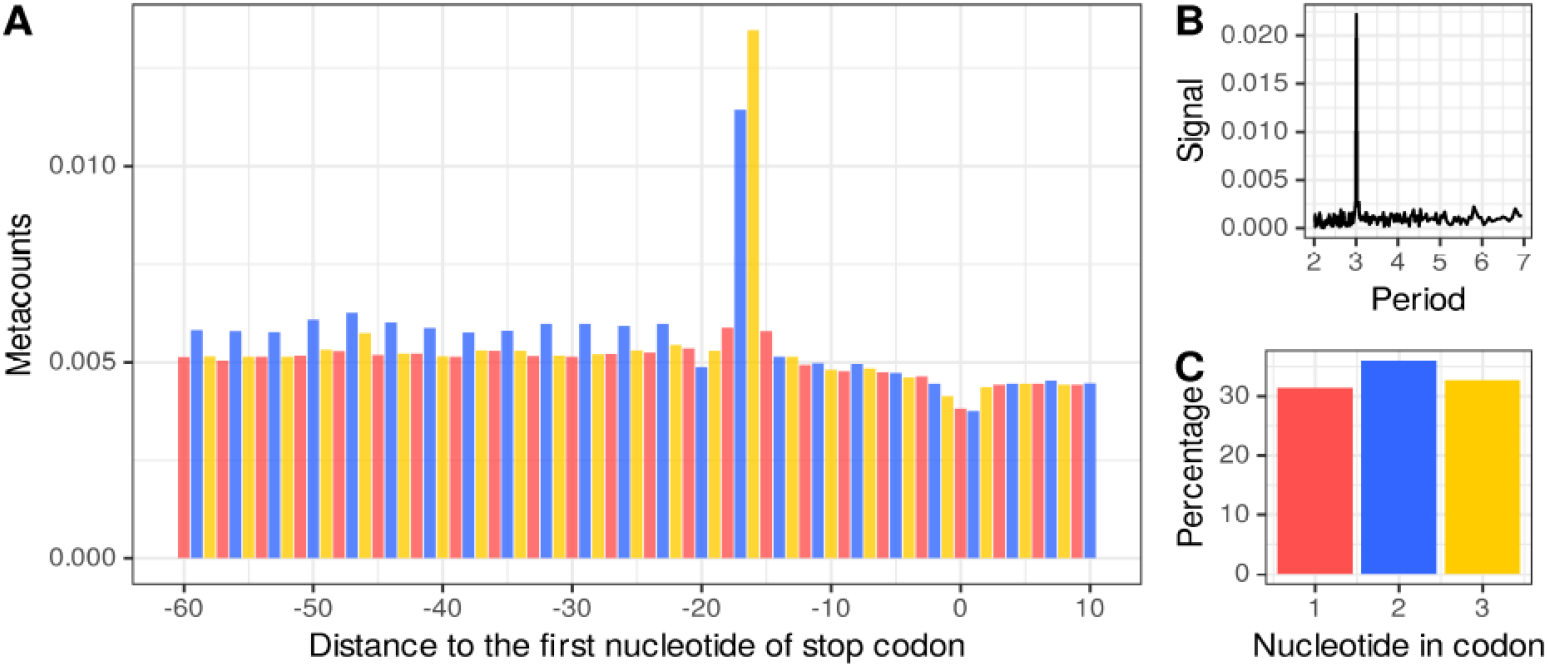
*A. thaliana* mRNA co-translational degradation by an exoRNase. (A) Metacount profile of *A. thaliana* CDS stop region. Each bar corresponds to the median number of normalized counts of 5’ ends mapped at relative distance to the first nucleotide of the stop codons. The first, second and third nucleotides of the codons in the translation frame, are indicated by red, blue and yellow bars, respectively. Counts are normalized by the total number of counts in the selected window. (B) Fast Fourier transform of a metacount profile obtained on a window size corresponding to the first quartile of the CDS length distribution. (C) Bar plot representing the percentage of normalized counts falling into each of the three nucleotides in the codons of all of the CDSs.

From these results, we can conclude that in *A. thaliana*, a part of the 5’ to 3’ exonucleolytic RNA degradation occurs simultaneously with the translation. This co-translational degradation of mRNA was demonstrated to be accomplished by the cytoplasm-localized 5’ to 3’ exoribonuclease XRN4 (Yu et al., 2016), resulting in a 3 nucleotide periodicity with significantly lower abundances in frame 1 corresponding to the boundary of the ribosome, which is consistent with the results obtained by *rnaends*.

The detailed analysis and code are available in a tutorial R notebook provided as supplementary material (see Data, scripts, code, and supplementary information availability section). The same type of analysis in *S. aureus* making use of EMOTE generated datasets in the study of RNase Y endoribonucleolytic cleavage (Khemici et al., 2015) is also provided in the supplementary material, where we show that there is a three nucleotide periodicity attributable to co-translational degradation, and a frame preference bias towards the second nucleotide in codons.

Here, we have focused on the stop codon as the feature of interest, however any other feature such as operons or unique position like TSSs can be used to centre the 5’ end distributions and obtain a metacount profile not necessarily related to the ribosome.

### Post-transcriptional modifications and differences in their proportions

At the other end of transcripts, *i.e*. at their 3’ end, RNAs can be exoribonucleolytically degraded, or non-template ribonucleotides can be added post-transcriptionally. The latter can, for example, be poly-adenylation of eukaryotic mRNAs, or the addition of CCA at the 3’ end of tRNAs that are transcribed from genes without CCA at the 3’ end of the genome template. In (Xu et al., 2023), a family of toxin-antitoxins present in *Mycobacterium tuberculosis* has been studied. One of the findings is that the toxins add ribonucleotides to the tRNA 3’ ends prior to amino acid loading onto the tRNA, thus preventing its use for translation by the ribosome.

Compared to the 5’ end analyses described so far, in which reads should align entirely and with 100% identity to the genome reference, the specificity for this type of analyses is that the reads are constituted by the end of the transcribed gene (up to 3’ end of the transcript) and possibly by an additional sequence that does not align to the reference sequence. To be able to align such reads, the *rnaends* package makes it possible to use *Rsubread* (Liao et al., 2019) as described in (Xu et al., 2023). In short, the reads are aligned to reference sequences allowing partial alignments (also termed soft-clipping). After mapping, only the locations of the 3’ end aligned regions are kept together with the unmapped regions downstream of the 3’ end. For quantification, the count table is obtained by grouping reads with identical 3’ locations and identical unmapped downstream sequences.

As an illustration of our implemented workflow, we used one of the raw FASTQ files published in (Xu et al., 2023) corresponding to *in vitro* transcribed tRNAs incubated or not with the MenT1 toxin with two replicates in each experimental condition. The workflow proceeds as follows. First, raw reads are validated and demultiplexed into FASTQ files for each condition and replicates, followed by alignment to reference tRNA sequences. Second, 3’ end locations are quantified taking into account UMIs (PCR amplification bias removal) and unmapped 3’ end downstream sequences. Third, mapped reads are filtered to keep only full length tRNAs possibly with 3’ end addition of cytidines (since *in vitro* transcribed tRNAs were incubated with the MenT1 toxin and cytidine triphosphate (CTP) (Xu et al., 2023)). A count table is then generated, which allow us to study the post-transcriptional modifications on tRNAs made by the MenT1 toxin. This is illustrated in Figure 4 where counts for each tRNA are represented by stacked bar plots. While in the control (without MenT1 treatment), tRNAs do not display 3’ end modifications, we can observe that there is a fraction of those having an additional C and, to a lesser extent, multiple Cs as shown in (Xu et al., 2023). Of note, CCC or CCCC additions are present in very low amounts for some tRNAs in some samples, and are negligible. Thus they are not visible in Figure 4, but are kept in the legend to indicate their existence.

**Figure 4.**
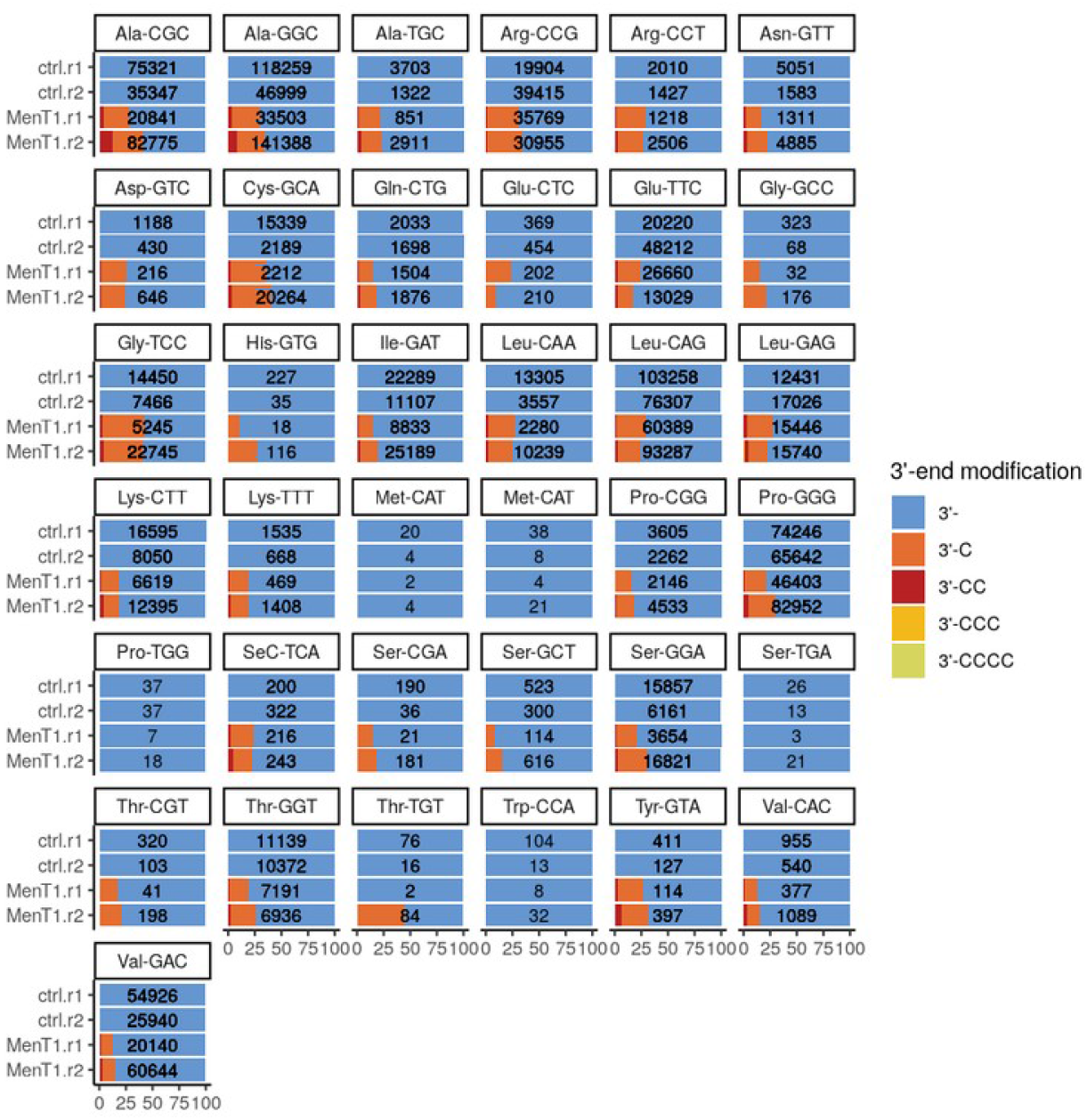
tRNAs 3’ ends modified by the MenT1 toxin were mapped by incubating *in vitro* transcribed tRNA from *Mycobacterium smegmatis* with or without MenT1 toxin in the presence of CTP. The total full length tRNAs (3’-) and full length with C additions are depicted with stacked bar plots for each tRNA represented as percentages of each full length tRNA with or without post-transcriptional additions. The two upper rows correspond to the two control replicates without MenT1 and the two lower rows correspond to the two samples treated with MenT1. The numbers on the bars indicate the number of mapped reads for each tRNA and sample. Only a few molecules with CCC or CCCC were detected, and are therefore not visible at the scale of the figure.

In addition to what was published in (Xu et al., 2023), *rnaends* can perform a statistical test to decide whether the proportion of native *vs*. modified tRNAs is significantly different in the two conditions and replicates (please note that for statistical analyses, more than two replicates is highly desirable). The statistical framework is different from differential gene expression which compares abundances that are discrete quantities. The goal in this analysis is to compare proportions based on abundances together with replicates. From all the tRNAs of Figure 4, only 5 do not have a significant difference in proportions: the two Met-CAT tRNAs, Pro-TGG, Ser-TGA and Trp-CCA tRNAs. This could be due to the very low counts observed for these tRNAs that would justify to remove them from the main findings. Apart from these five tRNAs, the results show that tRNAs exhibit similar levels of post-transcriptional additions, and that the MenT1 toxin does not seem to have preference for specific tRNAs, at least *in vitro*.

The workflow described here illustrates how to analyse only a subset of the data related to the MenT1 toxin generated by (Xu et al., 2023). Its adaptation to the processing of data generated for the other toxins investigated in this publication should be straightforward.

The detailed analysis and code are available in an R notebook provided as supplementary material (see Data, scripts, code, and supplementary information availability section).

## Discussion

In this article, we illustrated some of the potential uses of functions implemented in the *rnaends* package through scenarios typical of RNA metabolism studies. Throughout its development, we aimed at making choices to render it as generic and user friendly as possible. The functions and functionalities have been implemented as independently as possible, so that they can be reused in the broadest variety of situations related to the processing of RNA-end sequencing reads.

Existing software generally offer some but not all of the functionalities of *rnaends*, or are available in other programming languages or as command line tools. For instance, *MOIRAI* (Hasegawa et al., 2014), *RECLU* (Ohmiya et al., 2014), *CAGEr* (Haberle et al., 2015), *CAGEfightR* (Thodberg et al., 2019), are specific to TSS identification in eukaryotes and to the CAGE protocol. For a more general protocol, *TSSAR* (Amman et al., 2014) is available as Perl scripts that generate R code to process sequencing reads that have already been mapped (SAM or BAM files). Like *TSSr* (Lu et al., 2021), it is designed for TSS identification from exact 5’ ends but they do not cover other aspects of exact RNA-end sequencing such as the 3’ end or the signal periodicity analysis. Python tools or shell scripts generally focus on specific steps of the analysis: *READemption* (Förstner et al., 2014) is written in Python, and enables reads pre-processing, mapping and differential expression analysis for ‘classic’ RNA-Seq libraries, *i.e*. not dedicated to the study of the exact 5’ or 3’ ends of transcripts. *PyRanges* (Stovner & Sætrom, 2020), *Pygenomics* (Tamazian et al., 2023) and *Bioframe* (Open2C et al., 2024) also provide Python modules and tools for the analysis of genomic ranges, whereas our package focuses on exact ends of transcripts. *BigSeqKit* (Piñeiro & Pichel, 2022) targets large volumes of FASTA/FASTQ files but does not perform post-alignment tasks. *UMI-tools* (Smith et al., 2017) implements some functionalities for UMI deduplication. However, it does not allow to perform statistics on the UMI usage distribution and does not perform demultiplexing when barcodes have been embedded in the oligo during library preparation. It should be noted that *QuasR* (Gaidatzis et al., 2015) is an R/Bioconductor package that allows reads pre-processing (but mainly adapter removal, such as 3’AAAA for example), alignment and quantification in a *qCount* object. This latter offers more complex features for genomic intervals, but does not offer the simplicity of considering only 5’ or 3’ end counts of transcripts. The *BRGenomics* package (DeBerardine, 2023), also available in Bioconductor, is dedicated to the steps following reads mapping and therefore does not offer the pre-processing functions available in *rnaends. fivepseq* (Nersisyan et al., 2020), an easy to use software written in Python, performs comprehensive analyses of 5’P degradome datasets with respect to translational features and allows the global and gene-specific translational frame-preference and codon-specific ribosome protection patterns. However, it is restricted to 5’ end analyses. Overall, although several tools exist, they do not offer the possibility of performing all the analyses from an R environment and/or do not allow to focus on the specific aspects of analysing RNA 5’ or 3’ end sequencing data, notably by the means of a simple count matrix representation of their single nucleotide locations on the reference sequences.

The core of the *rnaends* package exploits the simplicity and versatility of the quantified sequencing reads. These reads can be represented as a table instead of genomic ranges. Indeed, for this type of studies, only the 5’ or 3’ end location on the reference sequences is of interest together with their abundance and strand, as well as potentially post-transcriptionally added ribonucleotides downstream their 3’ end. Efforts have been made to rely only on packages where necessary, and thus to only integrate functions from other packages into *rnaends* that already provide the required existing functionality. We hope these choices will allow non-experts to be able to pre-process raw read FASTQ files and map the reads to their reference sequences to obtain the count tables corresponding to their experiment.

In particular, the *rnaends* package provides the *read_features* table to describe the structure of the sequencing reads prior to their parsing, validation and optionally demultiplexing into separate FASTQ files. The few cases and patterns identified and implemented should provide various possibilities: from simple validation and counting of the raw reads from a FASTQ file, to the demultiplexing into different FASTQ files on the basis of a barcode integrated in the reads during the library preparation. It can also be used to trim reads or extract specific features such as UMIs.

Once the RNA-end count table has been obtained from the mapped reads, further downstream statistical analyses can be envisaged using the wide range of tests offered in R. This is illustrated in an R notebook for TSS identification in the on-line package documentation. Moreover, if required for analysis purpose, we have also developed dedicated R functions in *rnaends* as described when investigating degradation intermediates in *A. thaliana* (computation of metacount profiles, periodicity signal analysis) to highlight that a part of the 5’ to 3’ exonucleolytic RNA degradation occurs co-translationally. However, for more in-depth analyses of ribosome profiling, such as quantification for di- or tri-nucleotides or even amino acids, we recommend the use of *fivepseq* (Nersisyan et al., 2020). Post-transcriptional 3’ end modifications are also simply represented in the count table by adding the various sequences observed downstream the mapped 3’ end into an additional column specific to this kind of study. One can then explore the diverse 3’ end modifications of RNAs. It is also possible to compute statistics as illustrated with the tRNA modified or not by the MenT1 toxin.

On the other hand, our package can interoperate with packages providing some more elaborate data analysis such as differential expression with *edgeR* (Robinson et al., 2010) or *DESeq2* (Love et al., 2014). We set up an example to illustrate the simplicity of this integration through the identification of the cleavage sites of the endoribonuclease Y (RNase Y) in *Streptococcus pyogenes* based on the study of (Broglia et al., 2020) in a dedicated R notebook in the on-line package documentation. In particular, the authors have performed 5’ end sequencing on a wild-type strain and an RNase Y deletion mutant strain. Using the raw data provided with their publication, we show that differential expression analysis can be conducted using the data table structure generated by our package.

While *rnaends* provides functions encompassing complete workflows from pre-processing to downstream analyses, it was designed to be modular and the pre-processing, the quantification and the downstream analysis steps are independent. For instance, it is possible for the users to (i) pre-process FASTQ files with the *rnaends* functions, (ii) perform the reads mapping with an aligner not integrated in *rnaends* such as STAR (Dobin et al., 2013) widely used with eukaryotes for obtaining the BAM files, and (iii) obtain the count table and perform the downstream analysis with *rnaends* functions.

The *rnaends* package is under continuous development and features will be added in the future. The code documentation and the R notebooks should allow experts as well as less experienced programmers to pre-process and analyse RNA-end sequencing libraries.

## Acknowledgements

We are thankful to Julien Prados (University of Geneva, Switzerland) who initiated the use of R for EMOTE data analysis and made his source code public, and for his helpful comments. We would like to thank Reviewers for taking the time and effort necessary to review the manuscript. We sincerely appreciate all valuable comments and suggestions, which helped to improve the quality of the manuscript. A preprint version of this article has been peer-reviewed and recommended by PCI Genomics (https://doi.org/10.24072/pci.genomics.100440).

## Data, scripts, code, and supplementary information availability

The *rnaends* package is available on GitLab at (https://rnaends.gitlab.io/rnaends) and its version 1.0.0 has been deposited on Zenodo at https://doi.org/10.5281/zenodo.15076101 (Caetano et al., 2025). All RNA-Seq datasets used in this article are available in public repositories: RNA-Seq dataset used for TSS identification from (Prados et al., 2016) are in the GEO repository with accession number GSE85110. RNA-Seq dataset used for RNase Y endoribonucleolytic cleavage sites identification from (Broglia et al., 2020) are in the SRA NCBI repository with accession number SRP149896. RNA-Seq dataset used in degradation intermediates analysis in *A. thaliana* from (Hou et al., 2016) are in GEO repository with accession number GSE77549. RNA-Seq dataset used in degradation intermediates analysis in *S. aureus* from (Khemici et al., 2015) are in the GEO repository with accession number GSE68811. RNA-Seq dataset used in 3’ end post-transcriptional modifications from (Xu et al., 2023) are in the European Nucleotide Archive (ENA) at EMBL-EBI under accession number PRJEB62085.

